# Repetitive Sulfur Dioxide Exposure in Mice Models Post-Deployment Respiratory Syndrome

**DOI:** 10.1101/2023.05.15.540867

**Authors:** Sergey S. Gutor, Rodrigo I. Salinas, David S. Nichols, Julia M. R. Bazzano, Wei Han, Jason J. Gokey, Georgii Vasiukov, James D. West, Dawn C. Newcomb, Anna E. Dikalova, Bradley W. Richmond, Sergey I. Dikalov, Timothy S. Blackwell, Vasiliy V. Polosukhin

## Abstract

Soldiers deployed to Iraq and Afghanistan have a higher prevalence of respiratory symptoms than non-deployed military personnel and some have been shown to have a constellation of findings on lung biopsy termed post-deployment respiratory syndrome (PDRS). Since many of the deployers in this cohort reported exposure to sulfur dioxide (SO_2_), we developed a model of repetitive exposure to SO_2_ in mice that phenocopies many aspects of PDRS, including adaptive immune activation, airway wall remodeling, and pulmonary vascular disease (PVD). Although abnormalities in small airways were not sufficient to alter lung mechanics, PVD was associated with the development of pulmonary hypertension and reduced exercise tolerance in SO_2_ exposed mice. Further, we used pharmacologic and genetic approaches to demonstrate a critical role for oxidative stress and isolevuglandins in mediating PVD in this model. In summary, our results indicate that repetitive SO_2_ exposure recapitulates many aspects of PDRS and that oxidative stress may mediate PVD in this model, which may be helpful for future mechanistic studies examining the relationship between inhaled irritants, PVD, and PDRS.

## INTRODUCTION

Close to 3.5 million U.S. military service members have been deployed to Southwest Asia and Afghanistan over the last 20 years (1) and studies have shown an increase in respiratory symptoms in this group compared to non-deployed soldiers, with symptoms including cough, shortness of breath, and reduced exercise tolerance (2–4). Despite thorough medical examination, ∼30% of Veterans with exertional dyspnea remain undiagnosed by standard clinical evaluation (5). In 2011, 38 of 49 lung biopsies in soldiers with unexplained post-deployment exertional dyspnea were reported to show constrictive bronchiolitis (CB) (6). In 2021, we performed a comprehensive histopathological evaluation of lung biopsy samples from 50 soldiers with unexplained exertional dyspnea and exercise intolerance and found complex pathological changes in the distal lungs that included CB-like small airway remodeling, hypertensive-type vascular pathology, lymphocytic inflammation, and fibrosis involving alveolar tissue and pleura. We proposed the term “post-deployment respiratory syndrome” (PDRS) as a descriptor for the presence of this multi-compartment pathology in the distal lung in Veterans with exertional dyspnea following deployment to Iraq or Afghanistan (7). Also, deployment-related respiratory disease (DRRD) was recently proposed by a consensus statements as a broad term to subsume a wide range of potential syndromes and conditions identified through noninvasive evaluation or when surgical lung biopsy reveals evidence of multicompartmental lung injury that may include CB (8). Currently, there is no reliable model to investigate underlying pathogenetic mechanisms or test potential therapies in PDRS.

We reasoned that inhalation exposure to 125 ppm of sulfur dioxide (SO_2_), an environmental toxin to which some affected soldiers were exposed during the Mishraq State Sulfur Mine fire in Iraq in 2003, could be an appropriate stimulus to establish a murine model of PDRS (6). SO_2_ is a common air pollutant that rapidly forms sulfurous acid and bisulfite upon inhalation, leading to the production of reactive oxygen species (ROS), including sulfur trioxide anion that induces lipid peroxidation (9–15). Then, arachidonic acid and polyunsaturated fatty acids yields highly reactive isolevuglandins (isoLGs) via direct lipid peroxidation (11, 16, 17). IsoLGs rapidly adduct to lysine residues, resulting in protein crosslinking and functional alteration (17). Specifically, isoLGs can inhibit the mitochondrial deacetylase sirtuin-3 (Sirt3), leading to inactivation of the key antioxidant superoxide dismutase (SOD2) due to hyperacetylation (18).

To develop a murine model of PDRS after SO_2_ exposure, we optimized SO_2_ exposure parameters and performed a comprehensive histological and physiological analysis. In addition to showing that this model recapitulates important pathological aspects of PDRS, we identified long-term physiological abnormalities, including reduced exercise capacity and development of pulmonary hypertension. In addition, we showed that SO_2_-induced ROS production contributes to the lung pathology identified in this model. Some of the results of these studies have been previously reported in the form of abstracts (19).

## MATERIALS AND METHODS

### Mouse models

In most of the experiments, wild type (WT) mice on a C57Bl/6J background were used. Additionally, to investigate whether inhibition of the mitochondrial deacetylase Sirt3 contributed to the vascular pathology in SO_2_ exposed mice, we used transgenic mice that constitutively overexpress Sirt3 (∼4-fold) in all tissues based on the EIIa-Cre (which targets Cre expression in the early mouse embryo; OX-Sirt3 mice). To evaluate whether scavenging of isoLGs could prevent SO_2_-associated pathology, we treated mice with 2-HOBA (2-hydroxybenzylamine acetate). 2-HOBA was provided by the Department of Clinical Pharmacology at Vanderbilt University Medical Center. 2-HOBA was dissolved in the drinking water (1g/L). Mice were treated with 2-HOBA starting 3 days before exposure until the last exposure day. Fresh solutions were prepared every 3 days prior to animal dosing. All mice were 8-10 weeks of age and both male and female in equal proportion were used. Mice were housed in standard microisolators in the Vanderbilt animal care facility under a 12-hour light/12-hour dark cycle in a temperature-controlled room (24°C) with free access to food and water. All mouse experiments were performed according to a protocol approved by the Institutional Animal Care and Use Committee at Vanderbilt University Medical Center. SO_2_ exposure was performed in the animal care facility using an InExpose inhalational exposure system (SCIREQ) approved by our IACUC and institutional biosafety committee for this use. The system was operated under a standard fume hood and ppm was controlled with GasBadge® Pro Single-Gas Monitor (Industrial Scientific).

### Histopathological evaluation

Animals had been euthanized with subsequent harvesting of lungs. Lungs were perfused through the RV with sterile PBS until white. Left lungs were inflated with 10% buffered formalin using a 25-cm pressure column, fixed overnight, processed, embedded in paraffin, and cut into 5 μm sections. Right lungs were used for protein, RNA, and(or) cell isolation. Tissue sections stained for hematoxylin and eosin (H&E), PicroSirius red, and Verhoeff-Van Gieson (VVG) were used for routine histological examination and histomorphometry. Epithelial damage and infiltration scores were determined by a semi-quantitative severity score (0 to 3) for inflammatory cell infiltration and alveolar epithelial injury (0 to 3) as published elsewhere (20). Collagen content was measured as the area containing a specific fluorescence signal within the total area of the airway subepithelium on PicroSirius red-stained sections using a red fluorescence microscope technique. To assess airway wall remodeling, total subepithelial area was measured for each distal airway on digital microscopic images (×20 objective) on PAS-stained sections and then normalized to the reticular basement membrane (RBM) length. The area of media was measured for each artery adjacent to distal airway on digital microscopic images (×40 objective) on VVG stained sections and then normalized to the length of inner elastic membrane length. Muscularization of distal intrapulmonary vessels were determined on double immunostained tissue sections with anti- CD31 (rat monoclonal, #DIA-310, Dianova) and anti-α-SMA (rabbit monoclonal, # ab124964 (EPR5368), Abcam) antibody in different sized vessels (vessel inner diameter 10-25, 25-50, and 50-100 μm). Morphometric measurements were made by two independent pathologists blinded to study group using Image-Pro Plus 7.0 software (Media Cybernetics, Silver Springs, MD) or ImageJ 1.8.0 (NIH, Bethesda, MD).

### Generation of single cell suspensions and flow cytometry

Lungs were removed, minced with scissors, and incubated for 45 minutes at 37 °C in high glucose Gibco™ DMEM containing collagenase XI (0.7 mg/mL; C7657, Sigma-Aldrich) and type IV bovine pancreatic DNase (30 μg/mL; D5025, Sigma-Aldrich). The tissue was further disrupted by gently pushing undigested pieces through a 70 μm nylon screen with the plunger from a plastic 5 ml syringe. Cells were then pelleted at 200 × g, incubated in 1 ml of ammonium-chloride-potassium (ACK) lysing buffer for 5 min at room temperature to remove red blood cells, washed with PBS containing 0.5% fetal bovine serum (FBS) and 2 mM EDTA, and filtered through a second 70 μm nylon screen. Antibodies for flowcytometry utilized in this study were combined into two panels. <u>Macrophages, neutrophils, monocytes, DCs</u> (all BioLegend): Live-Dead Brilliant Violet 450 (Invitrogen, #L34955A), CD45-eFlour605 (#103140), CD11b-PE-Cy7 (#101216), CD11c-PE (#117308), Ly6G-APC-Cy7 (#127623), MHCII-FITC (#107606), CD64-APC (#139305), and F4/80-AF700 (#123130). <u>Lymphocytes</u> (all BioLegend): Live-Dead Brilliant Violet 450 (Invitrogen, #L34955A), CD45-eFlour605 (#103140), CD3-AF700 (#100216), CD4-PE-Cy5 (#100409), CD8-APC-Cy7 (#100714), NK1.1-PE-Cy7 (#108713), CD19-FITC (#115505), CD103-PE (#121405). The gating strategy for identifying DC subsets has previously been published (21). Flow cytometry was performed on various instruments in the Flow Cytometry Shared Resource Center and at the University of Florida. All analyses were performed using FlowJo software.

### Superoxide measurements using HPLC

Mouse lung 1 mm thick sections and isolated from mouse lung endothelial cells were loaded with DHE (50 µM) in KHB buffer by 30-minute incubation in a tissue culture incubator at 37 °C. Next, lung sections and endothelial cells were placed in methanol (300 µl) and homogenized with a glass pestle. The tissue homogenate was passed through a 0.22 μm syringe filter and methanol filtrates were analyzed by HPLC according to previously published protocols (22). The superoxide specific product 2-hydroxyethidium was detected using a C-18 reverse-phase column (Nucleosil 250 to 4.5 mm) and a mobile phase containing 0.1% trifluoroacetic acid and an acetonitrile gradient (from 37% to 47%) at a flow rate of 0.5 ml/min. 2-Hydroxyethidium was quantified by fluorescence detector using an emission wavelength of 580 nm and an excitation of 480 nm as described previously (23).

### Western blots

Endothelial cells were isolated from single cell suspension using LS Columns (#130-042-401, Miltenyi Biotec) and MACs magnetic beads with negative selection of CD45 (#130-052-301, Miltenyi Biotec), followed by positive selection of PECAM1/CD31 (#130-097-418, Miltenyi Biotec). Whole-cell lysates were prepared using RIPA buffer supplemented with protease (P8340, Sigma-Aldrich) and phosphatase inhibitors (P0044, P5726, Sigma-Aldrich), nicotinamide (98-92-0, Millipore Sigma), and TA (58880-19-16, Millipore Sigma). Protein concentrations were determined by bicinchoninic acid (BCA) assay and 15 μg of protein was separated on a 10% acrylamide gel and transferred to a nitrocellulose membrane.

Immunoblotting was performed using antibodies against isoLGs (D11), SOD2 (Santa Cruz Biotechnology, sc-30080), acSOD2(Abcam, ab137037), Sirt3(Cell Signaling, #5490), and GAPDH (Abcam, ab8245). Densitometry was performed using ImageJ 1.8.0 (NIH, Bethesda, MD).

### Hemodynamic measurements

Invasive hemodynamic measurements were conducted according to the previously described protocol (24). In brief, mice were given 0.75 mg/g of 2.5% avertin (a mixture of tert-amyl alcohol and 2,2,2-tribromoethanol; Sigma-Aldrich) to induce anesthesia. Then mice had been placed on a heating pad, and the jugular vein was dissected and catheterized with 1.4-F Mikro-Tip catheter (Millar Instruments, Houston, TX). The RV pressure tracing was then recorded utilizing Chart 5.3 software (AD Instruments). Immediately following invasive hemodynamic measurements, animals were euthanized with a subsequent harvest of lungs and hearts. The heart was excised with removal of the atria, and the RV and left ventricle (LV) plus septum were isolated for the measurement of the Fulton index (RV∶(LV + S)) as recommended (25).

### Exercise capacity

To estimate maximal exercise capacity, mice were placed in an enclosed single-lane treadmill (Columbus Instruments, Columbus, OH). The treadmill had 0 degrees incline during the whole experiment. At the beginning of the experiment, the mice started running on a treadmill at a speed of 10 m/min following a 10-min sedentary period. When the mouse reached the top of the belt, it was considered to have acclimated to that speed, and the speed was increased by 4 m/min every 3 min until the mouse reached exhaustion. Exhaustion was defined as >5 s time period when mice stop running and remained on the shock grid at the back of the treadmill and physical signs of exhaustion such as breathlessness and postural disturbances. The electrical stimulus was set at 163 volts with a stimulus rate of 4 pulses/sec and the amperage was set at 1.0 milliamps based upon manufacturers suggestions. If fatigue was observed, then mice were returned to their cages. For the acclimations (two occasions, at least 48h prior to the actual stress test) mice were initiated running at 10 m/min (0% incline) for 10 minutes following a 10 min sedentary period.

### Pulmonary function testing

Lung function measurements were quantified using the FlexiVent apparatus (SCIREQ). Mice were anesthetized with pentobarbital sodium (85 mg/kg), and an 18-gauge tracheostomy tube was placed in the trachea. Mice were then mechanically ventilated using the SCIREQ FlexiVent apparatus with 150 breaths/min and a tidal volume of 10 mL/kg body weight prior to lung function measurements. Respiratory system resistance, elastance and compliance were captured using the FlexiVent “Snapshot model”.

### Statistical analysis

The normality of continuous variable distribution was examined with the Shapiro-Wilk test. Comparisons of the normally distributed continuous variables between two groups were assessed by the t-test and more than two groups by One-way ANOVA. Otherwise, we used appropriate non-parametric tests (Mann-Whitney and Kruskal-Wallis for 2 and >2 group comparisons, respectively). For multiple comparisons, we conducted pairwise comparisons among study groups using the Student’s t-test or Mann-Whitney U test and p-values were Bonferroni adjusted for multiple comparisons. Adjusted p-values less than 0.05 were considered statistically significant. χ2 test or Fisher’s exact test was used for categorical variables. All analyses were performed using R-software version 3.5.2 (www.r-project.org).

## RESULTS

### SO_2_ exposure leads to airway wall remodeling, inflammation, and vascular pathology

For model development, we focused on epithelial injury and airway wall remodeling (collagen deposition and wall thickening) as primary end points. We first performed experiments to optimize the duration of exposure (hours) using a moderate SO_2_ concentration (125 ppm) for 2 or 4 hours. At 24 hours post-treatment, all mice exposed to SO_2_ showed evidence of epithelial injury and impaired epithelial barrier function. Using a previously described semi-quantitative scoring system (20), we showed that 4 hours of SO_2_ exposure caused more severe epithelial damage compared to 2 hours of exposure (Figure 1B). Pathological findings included epithelial cell desquamation with epithelial cell clumps and agglomerates in airway lumen, along with areas of the denuded reticular basal membrane (Figure 1A). Next, we studied the time course for airway remodeling after a single exposure to 125 ppm SO_2_ for 4 hours by measuring collagen content within the subepithelium of small airways on PicroSirius red-stained sections at 36 hours, 1, 2, 4, and 8 weeks after exposure. Collagen deposition increased progressively after exposure, peaking at 4 weeks post-treatment (Figure 1C,D). After choosing a dose, exposure duration, and post-treatment timepoint, we tested whether multiple SO_2_ exposures could further amplify small airway remodeling. We tested three dosing strategies: 1) a single 4-hour exposure to 125 ppm SO_2_ (x1), 2) weekly exposure to 125 ppm SO_2_ (4 hours duration) for 3 consecutive weeks (x3), and 3) five daily exposures to 125 ppm SO_2_ (4 hours duration) for 2 consecutive weeks (x10). At 4 weeks after the last SO_2_ exposure, we found that subepithelial thickness was maximal in small airways from the group treated with 10 total exposures to SO_2_ (Figure 1E,F).

**Figure 1.**
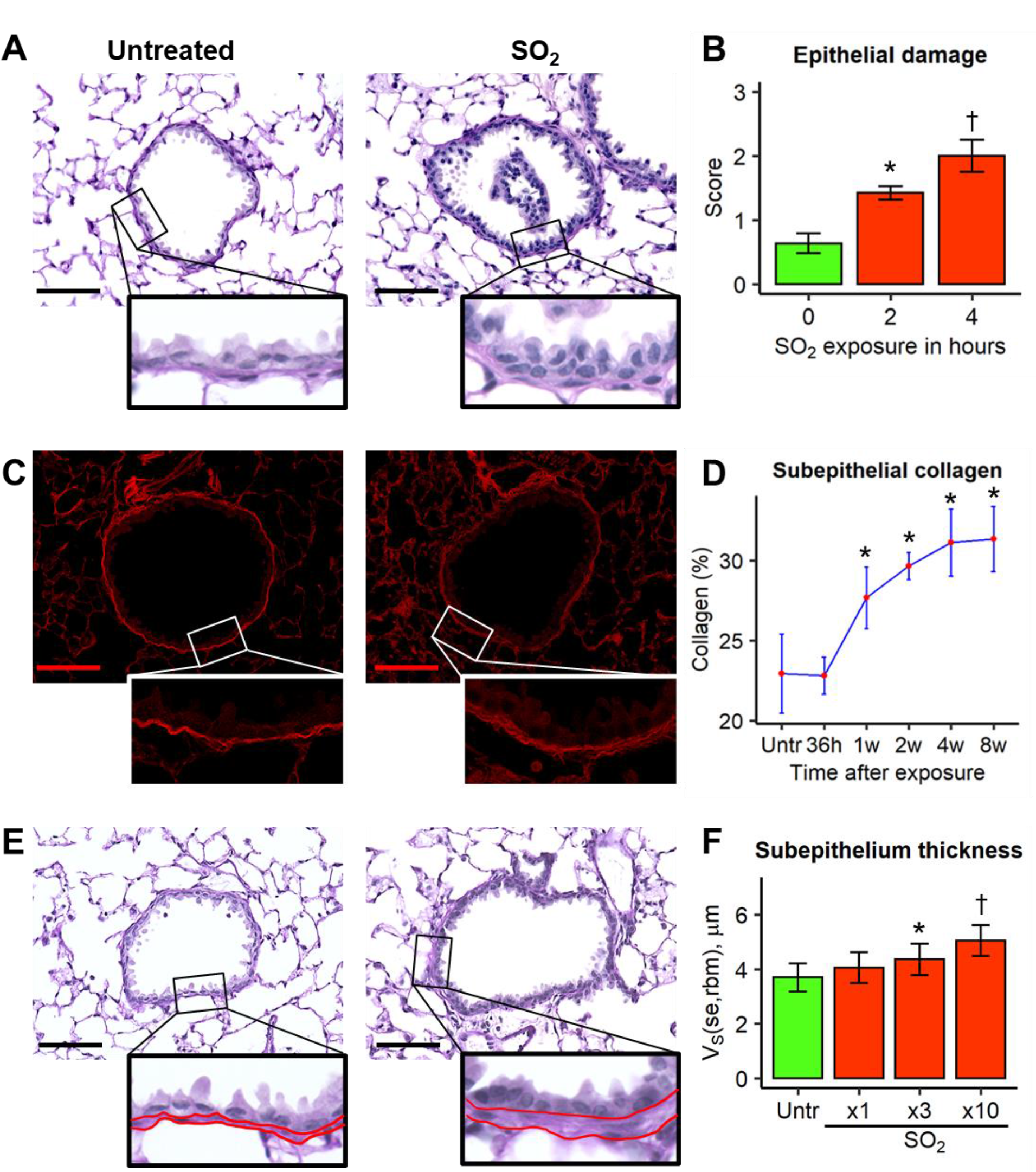
SO_2_ exposure model optimization and validation. A, normal-appearing distal airway in untreated WT mice (left) and epithelial sloughing and epithelial cell agglomerate in distal airway lumen of 125 ppm SO_2_-exposed mice for 4 hours (right). Periodic acid-Schiff (PAS) staining. Scale bars = 50 µm. B, epithelial damage score after two and four hours of 125 ppm SO_2_ exposure. Groups were compared using Student’s t-test and p-values were Bonferroni- adjusted for multiple group comparisons. * p<0.05 compared to untreated mice, † p<0.05 compared to untreated and SO_2_-exposed for 2 hours mice. N = 4 mice per group. C, normal- appearing distal airway in untreated WT mice (left) and collagen deposition in distal airway wall of 125 ppm SO_2_-exposed mice for 4 hours at 4 weeks after the last exposure (right). PicroSirius red staining. Scale bars = 50 µm. D, collagen content in distal airway wall increased exponentially over post-exposure time. Data presented as mean (red dots) ± SD (whiskers). Groups were compared using Student’s t-test and p-values were Bonferroni-adjusted for multiple group comparisons. * p<0.05 compared to untreated mice. N = 3-12 mice per time point. E, normal-appearing distal airway in untreated WT mice (left) and distal airway wall thickening after 10 days of 125 ppm SO_2_-exposed mice for 4 hours and 4 weeks of post-exposure time (right). Red lines outline the subepithelial layer. PAS staining. Scale bars = 50 µm. F, dose- dependent distal airway wall thickening. V_S_(se,rbm) – subepithelium thickness. Groups were compared using Student’s t-test and p-values were Bonferroni-adjusted for multiple group comparisons. * p<0.05 compared to untreated mice, † p<0.05 compared to untreated, single and three times SO_2_-exposed mice. N = 12 mice per study group.

Based on the findings described above, we chose the x10 model for subsequent studies (Figure 2A). Consistent with the increase in subepithelial thickness in the model, we identified prominent collagen deposition in distal airway walls compared to untreated mice (Figure 2B,C). At 4 weeks after the last SO_2_ exposure, we isolated lung cells and performed flow cytometry analysis for inflammatory cells using a previously published gating strategy (21). T lymphocytes (including CD3+, CD4+, and CD8+ cells), type 1 and type 2 conventional and monocyte-derived dendritic cells (cDC1, cDC2, and moDC), and lung macrophages were increased in SO_2_-exposed mice compared to controls (Figure 2D-F). Neutrophils, however, were similar between SO_2_- treated and control mice (Figure 2F). These findings are consistent with our previous report of persistent adaptive immune activation in the lungs of soldiers with PDRS (26).

**Figure 2.**
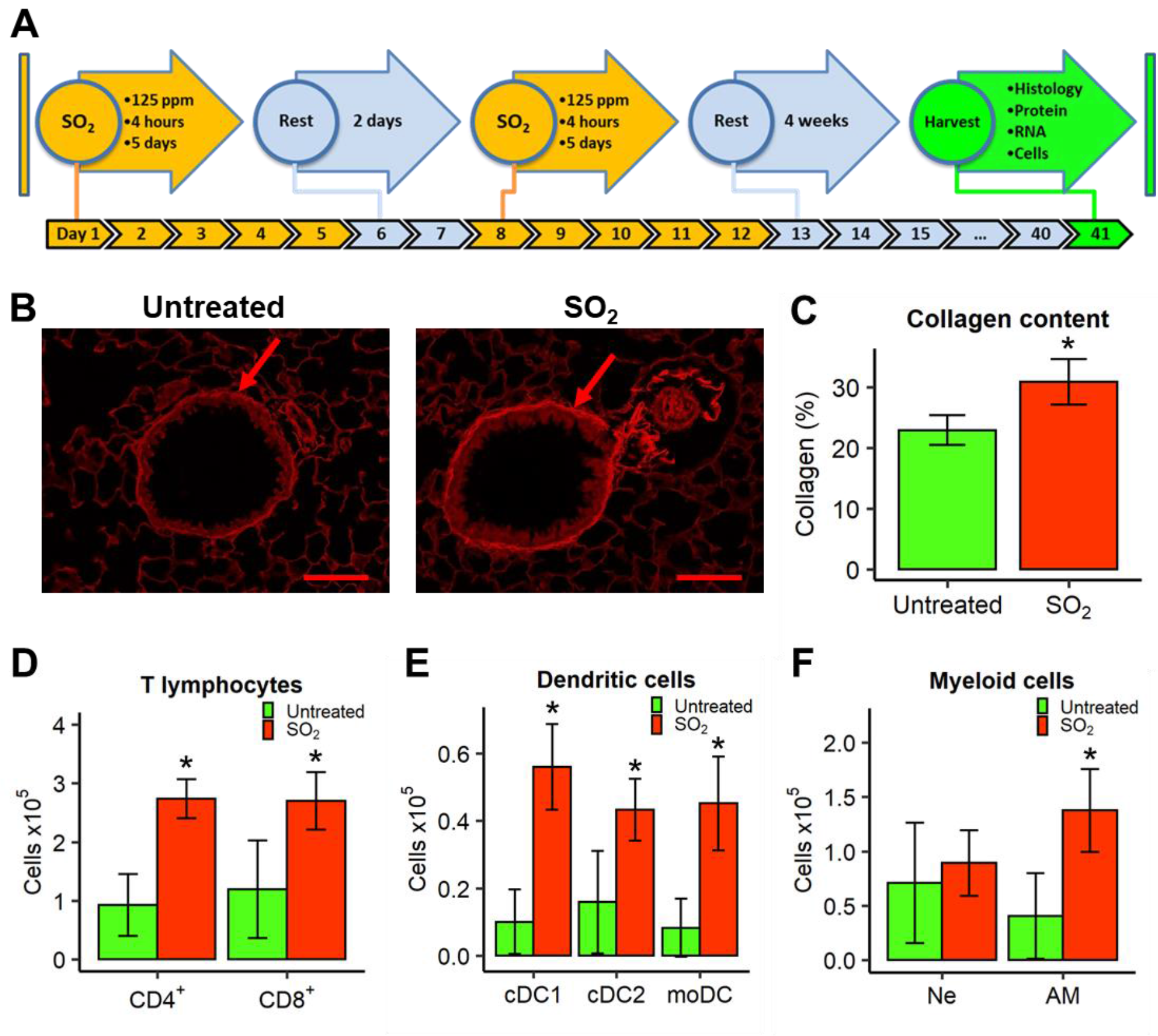
SO_2_ inhalation exposure leads to distal airway remodeling and chronic inflammatory cell influx. A, strategy and timeline of SO_2_ exposure for reproduction of PDRS- associated pathological lesions in mice. B, normal-appearing bronchovascular bundle in untreated WT mice; increased collagen deposition in distal small airways in WT mice exposed to 125 ppm SO_2_ for 4 hours a day, 5 days per week for 2 weeks, followed by 28 days post-exposure period (red arrow). PicroSirius red staining. Scale bars = 50 µm. C, collagen deposition in distal airways. Groups were compared using Student’s t-test. * p<0.001 compared to untreated mice. N = 12 mice per study group. D-F, increased numbers of CD4+ and CD8+ T cells, conventional type 1 (cDC1) and 2 (cDC2) and monocyte-derived (moDC) dendritic cells, and alveolar macrophages (AM) in SO_2_-exposed mice compared to unexposed age-matched mice. Ne – neutrophils. Groups were compared using Student’s t-test. * p<0.01 compared to untreated mice. N = 6 mice per study group.

To further investigate the spectrum of pathological abnormalities in our model, we performed histological evaluation of the pulmonary vasculature in mice exposed to vehicle or SO_2_ (x10 model). We found increased medial thickness in pulmonary arteries adjacent to small airways (Figure 3A,B). In addition, we detected increased muscularization of distal intra-acinar microvasculature (vessels with diameter less than 50 μm) in mice exposed to SO_2_ compared to controls (Figure 3C). Together, these studies showed that this model recapitulates key features of PDRS, including persistent airway remodeling, chronic lymphocytic inflammation, and vascular pathology.

**Figure 3.**
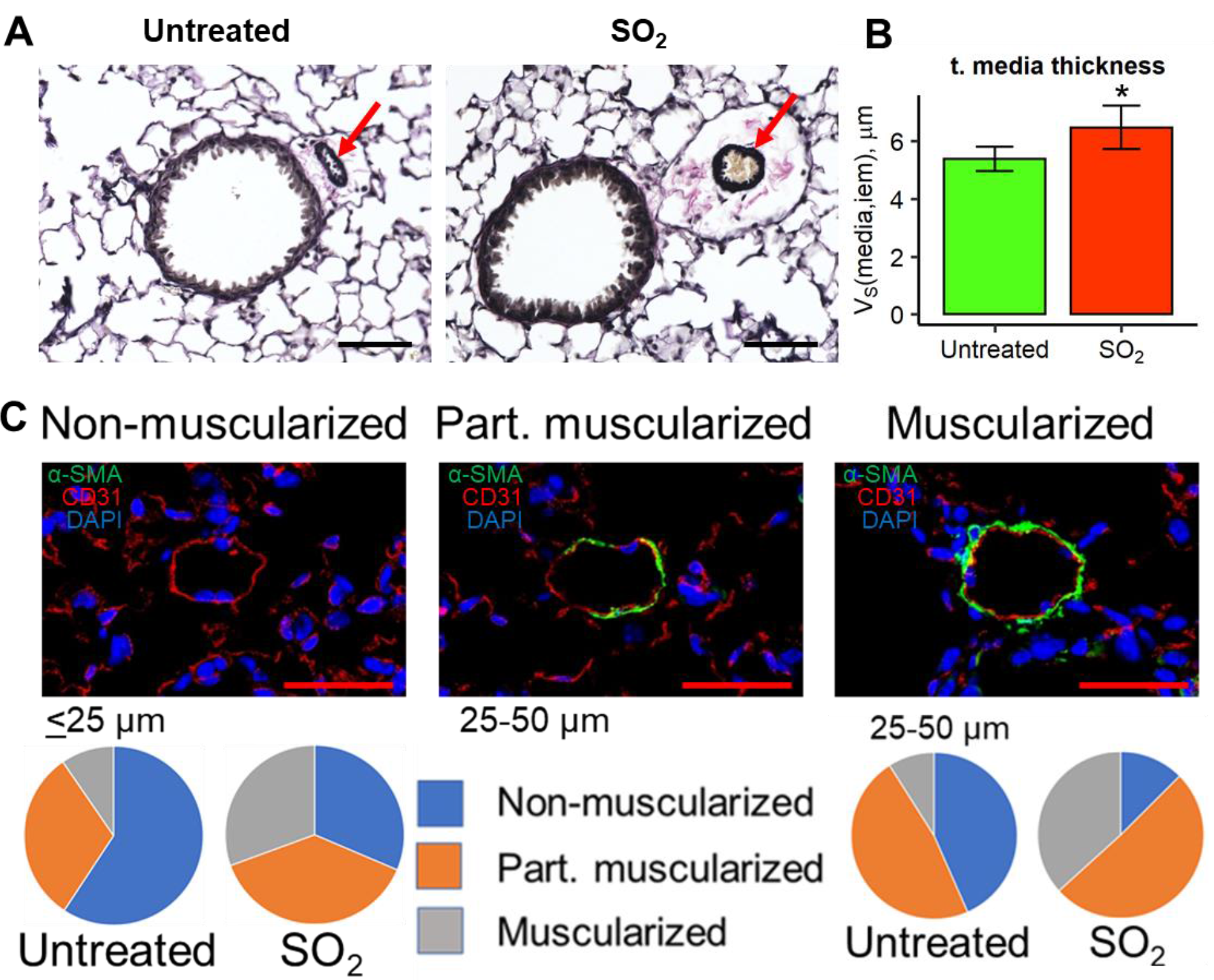
Effects of SO_2_ inhalation exposure on pulmonary vasculature. A, normal-appearing bronchovascular bundle in untreated WT mice; increased medial thickness (red arrow) and fibrosis of adventitia in pulmonary artery in WT mice exposed to SO_2_. Verhoeff-Van Gieson staining. Scale bars = 50 µm. B, bar plot showing *tunica media* thickness, V_S_(media,iem), in adjacent to distal airway pulmonary arteries. Groups were compared using Student’s t-test. * p<0.001 compared to untreated mice. N = 12 mice per study group. C, images illustrate examples of non-muscularized, partially muscularized (<50% vessel circumference with smooth muscle) and muscularized (>50% vessel circumference with smooth muscle) blood vessels with size <50 µm. Double immunofluorescence with anti-CD31 (red) and anti-α-smooth muscle actin (green) antibodies. Scale bars = 25 µm. 2D Pie charts illustrate the distribution of non-muscularized, partially muscularized and muscularized intra-acinar blood vessels in untreated and SO_2_ exposed to mice. p < 0.01 for group exposed to SO_2_ compared to untreated control mice by Chi-square test. N = 12 mice per study group.

### Repetitive SO_2_ exposure induce pulmonary hypertension with reduced exercise capacity

To examine whether pathological abnormalities observed in the lungs of mice exposed to SO_2_ resulted to persistent effects on cardiorespiratory function, mice were subjected to FlexiVent-based lung function analysis at 6 months after completion of the SO_2_ exposure protocol (x10 model). No differences in lung function were identified, including inspiratory capacity, total respiratory system resistance, compliance, and elastance (Figure 4A-C). We also examined exercise capacity in mice following SO_2_ exposure using incremental treadmill testing. Initial treadmill speed was set at 10 m/min (step 1) and increased every 3 min by 4 m/min until mice reached exhaustion. We found that SO_2_-exposed mice (x10 model) had significantly reduced exercise capacity compared to control (untreated) mice (Figure 4D).

**Figure 4.**
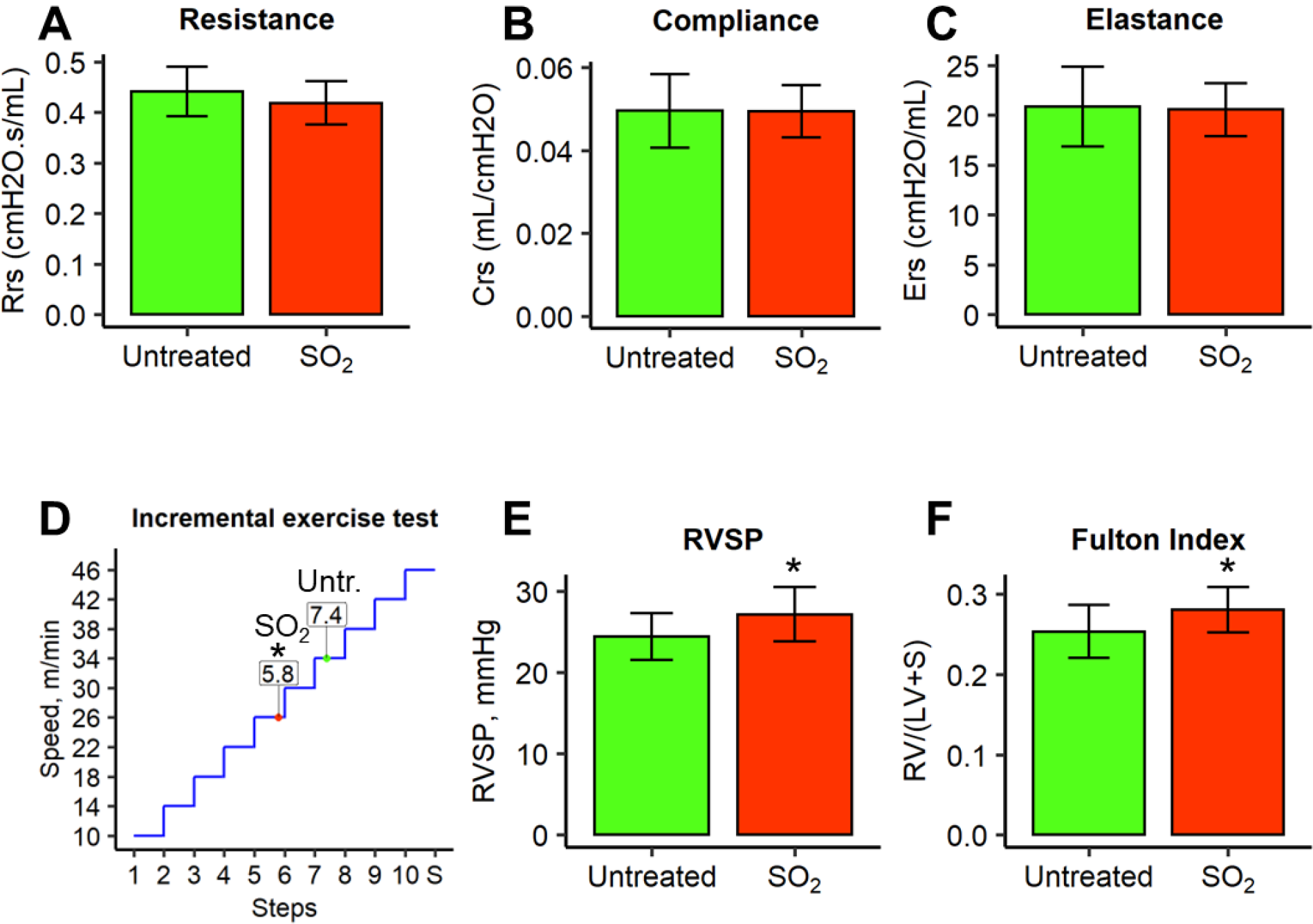
Long-term physiological effects of SO_2_ exposure. A-C, total respiratory system resistance (A), compliance (B), and elastance (C) measurements by the FlexiVent “Snapshot model”. Groups were compared using Student’s t-test. N = 11-12 mice per study group. D, incremental treadmill exercise test showing reduced maximal running speed in SO_2_ exposed mice (red) compared to unexposed mice (green). * p<0.05 compared to untreated mice (Student’s t-test). E-F, increase of RV systolic pressure (RVSP) and Fulton index in SO_2_- exposed mice compared to unexposed mice. Pulmonary hemodynamics were analyzed via the jugular vein using 1.4-F Mikro-Tip catheter. All experiments are conducted 6 months after the last exposure. Groups were compared using Student’s t-test. * p<0.05 compared to untreated mice. N = 17-18 mice per study group.

We then questioned whether the vascular lesions associated with SO_2_ exposure could result in pulmonary hypertension (PH). In another set of mice, we performed pressure tracings with a catheter placed in right ventricle (RV) through the jugular vein. RV systolic pressure (RVSP) was measured followed by measurement of the Fulton index [RV/(LV+S)] on excised hearts. These studies revealed increased RVSP at 6 months after SO_2_ exposure compared to controls (Figure 4E) as well as increase in Fulton index (Figure 4F). These studies identified long-term physiological consequence of SO_2_ exposure in our model, including exercise limitation, PH, and RV hypertrophy.

### SO_2_ exposure leads to chronic superoxide production with formation of isolevuglandins

Since it has been reported that exposure to SO_2_ induces oxidative stress in the lungs, we performed DHE-based assessment of superoxide production by HPLC analysis on thin-cut (1 mm thick) sections of a left lung and endothelial cells isolated from lung tissue. We showed increased 2OH-ethidium (a specific product of superoxide) normalized to protein content in lung tissue (Figure 5A), consistent with induction of oxidative stress by SO_2_. In addition, we isolated pulmonary endothelial cells from these mice and found a 2-fold increase in 2OH-ethidium levels (Figure 5B,C) following acute SO_2_ exposure.

**Figure 5.**
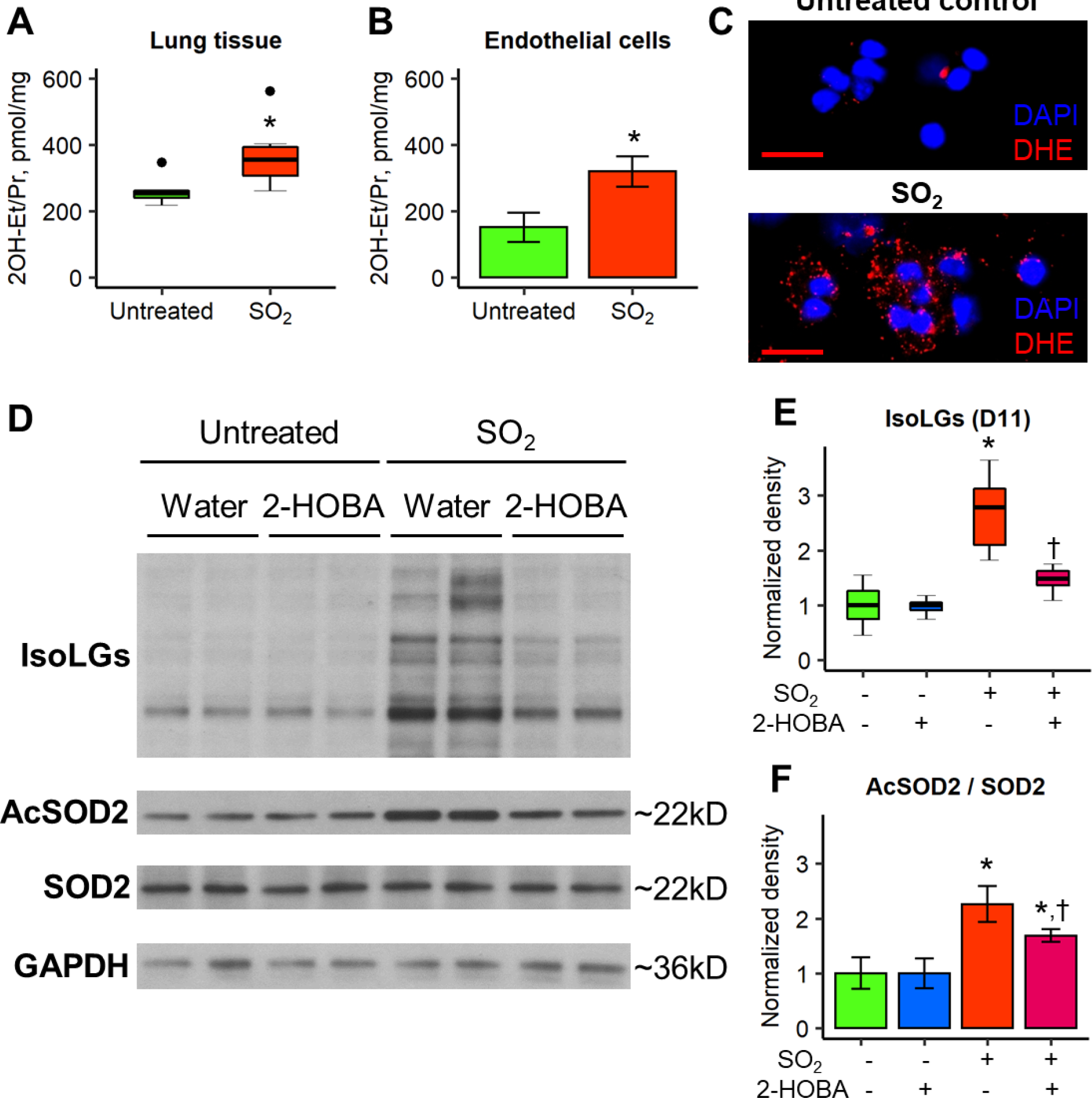
ROS and IsoLGs accumulation with downstream inactivation of Sirt3 and SOD2 after SO_2_ exposure. A-B, plots showing increased 2OH-ethidium (marker of superoxide) normalized to protein content (2OH-Et/Pr) in lung tissue 1 mm thick section (A) and endothelial cells isolated from lungs (B). Groups were compared using Mann-Whitney U-test (A) or Student’s t-test (B). N = 4-6 mice per study group. C, ROS accumulation in endothelial cells after SO_2_ exposure compared to vehicle. Dihydroethidium (DHE) stained in red. Scale bars = 25 µm. D, representative western blots for proteins with isoLG adducts and SOD2 acetylation in isolated endothelial cells. Endothelial cells were isolated from lungs of untreated or SO_2_-exposed mice with or without supplementation of isoLG scavenger 2-HOBA (1 g/L in drinking water). E and F, densitometry graphs showing accumulation of isoLG-protein adducts (E) and SOD2 hyperacetylation (F) in SO2-exposed mice and effect mitigation by 2-HOBA supplementation. Groups were compared using Mann-Whitney U test (E) or Student’s t-test (F) and p-values were Bonferroni-adjusted for multiple group comparisons. * p<0.05 compared to unexposed mice, † p<0.05 compared to SO_2_-exposed mice. N = 6 mice per study group.

Next, we wondered whether chronic ROS production was present in our repetitive SO_2_ exposure model and was linked to the persistent pathology. The result can be a positive feedback of ROS production and accumulation of isoLG-adducted proteins. To assess this pathway in our model, we measured isoLG-adducts by western blot analysis using the D11 antibody that recognizes isoLG-lysine adducts (27) and antibodies against SOD2 and acetylated SOD2. We found an accumulation of isoLG-adducted proteins (Figure 5D,E) as well as hyperacetylation of SOD2 (Figure 5D,F), consistent with Sirt3 inactivation.

To evaluate whether scavenging of isoLGs could protect from SOD2 hyperacetylation, we treated mice with the isoLG scavenger 2-Hydroxybenzylamine (2-HOBA) in drinking water (1g/L) starting 3 days prior to initiation of the SO_2_ (x10 model) and continued throughout the exposure period. At 4 weeks after the last SO_2_ exposure, we found that 2-HOBA treatment reduced formation of isoLG adducts and reduced SOD2 acetylation in endothelial cells (Figure 5D-F), thereby suggesting that isoLGs are responsible for Sirt3 inactivation in our model. These findings suggest that exposure to SO_2_ causes chronic ROS production, accumulation of isoLG-adducted proteins, and SOD2 hyperacetylation in pulmonary endothelial cells.

### 2-HOBA and Sirt3 overexpression limit vascular remodeling after SO2 exposure

We postulated that chronic ROS production and isoLG formation could drive the vascular pathology in our model of repetitive SO_2_ exposure through Sirt3 inactivation. Therefore, we treated mice with the isoLG scavenger 2-HOBA in the drinking water (1g/L) as described above and performed histological analysis of lung tissue collected from these mice 4 weeks after completion of the exposure protocol. Although we found no significant impact of 2-HOBA on collagen deposition (Figure 6A) or subepithelial thickness (Figure 6B) in distal airway walls, 2- HOBA treatment prevented of *tunica media* thickening in pulmonary arteries adjacent to small airways (Figure 6C) and muscularization of distal intra-acinar microvasculature (Figure 6D) in SO_2_-exposed mice.

**Figure 6.**
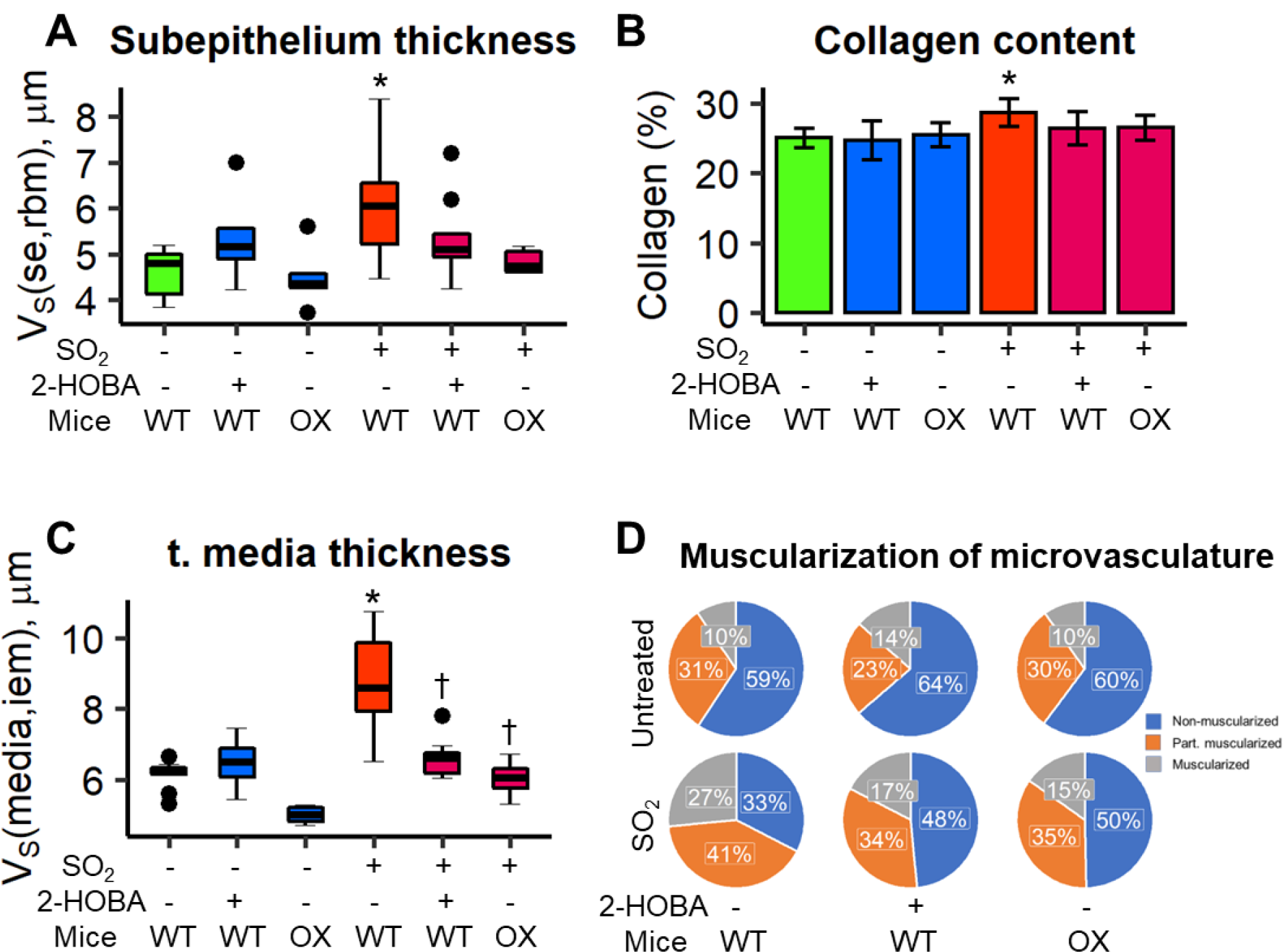
2-HOBA supplementation and Sirt3 overexpression prevent vascular remodeling after SO_2_ exposure. A and B, graphs showing distal airway wall thickening (A) and collagen deposition (B) in exposed to SO_2_ WT or Sirt3 overexpressing (OX) mice with or without 2- HOBA supplementation. V_S_(se,rbm) – subepithelium thickness. C, graph shows protective effect of 2-HOBA and Sirt3 overexpression on increased medial thickness in pulmonary arterioles adjacent to distal airways in WT mice exposed to SO_2_. V_S_(media,iem) – *tunica media* thickness. Groups were compared using Mann-Whitney U test (A,C) or Student’s t-test (B) and p-values were Bonferroni-adjusted for multiple group comparisons. * p<0.05 compared to untreated mice, † p<0.05 compared to SO_2_-exposed mice. N = 5-11 mice per study group. D, 2D Pie charts illustrate distribution of non-muscularized, partially muscularized and muscularized intra-acinar blood vessels (<25 μm in size) in WT and Sirt3 OX untreated or repetitive exposed to SO_2_ mice with or without 2-HOBA supplementation. p < 0.01 for group exposed to SO_2_ compared to untreated control mice by Chi-square test. N = 6-12 mice per study group.

To investigate whether inhibition of the mitochondrial deacetylase Sirt3 contributes to the vascular pathology in SO_2_ exposed mice, we utilized transgenic mice that constitutively overexpress Sirt3 (∼4-fold) in all tissues based on the EIIa-Cre (which targets Cre expression in the early mouse embryo) (28). Sirt3 overexpressing (OX) and wild-type (WT) mice were exposed to SO_2_ (x10 model) or remained untreated, and lung tissue was collected 4 weeks after completion of the protocol. We found no differences in collagen deposition (Figure 6E) or subepithelial thickness (Figure 6F) in distal airway walls of Sirt3 OX mice compared to WT. However, Sirt3 OX mice exhibited a significant reduction in *tunica media* thickness in pulmonary arteries adjacent to small airways, reduced *tunica media* thickening in arteries adjacent to small airways (Figure 6G), and a reduction of neomuscularization of microvasculature (Figure 6H). Cumulatively, these data indicate that SO_2_-induced vascular remodeling is driven, at least in part, by isoLG accumulation and downstream inactivation of Sirt3.

## DISCUSSION

In our model, we found that repetitive SO_2_ exposure models epithelial injury followed by airway wall remodeling, including subepithelial thickening and collagen deposition, chronic inflammatory cell influx, and pulmonary vascular remodeling, which are consistent with pathological lesions and persistent adaptive immune activation previously reported in lungs of Veterans with PDRS (7, 26, 29). Further analysis showed a long-term physiological consequence of SO_2_ exposure that overlaps with clinical symptoms of Veterans with PDRS, including exercise limitation without significant changes in lung function. Furthermore, our studies revealed that long-term effects of SO_2_ exposure include RV hypertrophy and development of PH. In mechanistic studies, we showed that SO_2_ exposure results in chronic ROS production and accumulation of isoLG-adducted proteins, inactivation of Sirt3, and SOD2 hyperacetylation in pulmonary endothelial cells, which impacts vascular remodeling in the distal lung. Thus, our model recapitulates pathological and physiological features of PDRS and suggests oxidative stress could be a molecular pathway underlying vascular pathology in this disease.

PDRS has been proposed as a descriptor for the combination of: 1) history of deployment in Southwest Asia and Afghanistan, 2) inhalational exposures during deployment, 3) chronic respiratory symptoms (reduced exercise tolerance and exertional dyspnea) that develop and persist in the post-deployment period, and 4) lung pathology that affects all distal lung compartments (7). These individuals typically have normal (or mildly abnormal) pulmonary function testing and CT scans of the lungs, making diagnosis challenging (6, 7). Pathological features of PDRS include fibrous remodeling of bronchiolar walls consistent with CB, with smooth muscle hypertrophy and increased collagen deposition. The disease also involves medial thickening (due to smooth muscle hyperplasia/hypertrophy) in distal pulmonary arteries adjacent to small airways, abnormal muscularization, excessive collagen deposition, and increased wall thickness in intra-acinar pulmonary arteries, as well as loss of capillaries in alveolar septa. These findings were first reported by King et. al. in 2011 and have been extended by our group and others (6, 7, 26, 29). Additionally, these lungs show evidence of an ongoing adaptive immune response (T cell activation) with diffuse infiltration of CD4 and CD8 T cells and B cell-containing lymphoid follicles (26). Together, available lung biopsy data indicate that PDRS is a complex lung disease involving multiple components of the distal lungs. Currently, there is limited information on the natural history of PDRS and the specific factors driving exertional dyspnea and exercise limitation in these patients remain incompletely understood.

Although we chose SO_2_ to model this disease because of the reported SO_2_ exposure of many soldiers in the initial cohort (6), the diverse effects of SO_2_ exposure on the respiratory system necessitate careful selection of model conditions to reproduce PDRS. One of the key aspects was SO_2_ concentration, since low concentrations of the gas (5-30 ppm) are reported to affect the upper respiratory tract only (30–33), while high concentrations (250-800 ppm) can cause severe damage and death related to sloughing of the bronchial epithelium (34–37). Thus, SO_2_ inhalation has been associated with respiratory injuries of varying severity depending on exposure conditions, such as airway epithelial injury, peribronchiolar inflammation and fibrosis, alveolar edema, and thickening of the alveolar-capillary barrier (9, 31, 34, 38–40). Consistent with prior reports, our study found that exposure to moderate concentration of SO_2_ (125 ppm) for 4 hours was sufficient to induce epithelial injury throughout the airways (31, 34, 41, 42), which was followed by excessive collagen deposition in airway wall at 4 weeks after exposure. By repetitive SO_2_ exposure, we were able to induce substantial distal airway wall remodeling and other pathological features of PDRS. Similarly, chronic (up to 45 days) exposure to 250-400 ppm SO_2_ for 4-5 hours a day has been used to model chronic obstructive disease, chronic bronchitis, and proliferative bronchiolitis (30, 40, 43, 44). Consequently, the variability in respiratory symptoms and diseases observed in Veterans, including asthma, airway hyperreactivity, and PDRS, may be attributed to the level of SO_2_ that were present at different deployment sites.

One of the key aspects of PDRS is chronic lymphocytic inflammation and our study has shown increased numbers of T lymphocytes, dendritic cells, and lung macrophages at 4 weeks post SO_2_ treatment. It is consistent with previous studies that demonstrated chronic exposure (20- 45 days) to SO_2_ causes macrophage, lymphocyte, and plasma cell infiltration in the lung tissue (40, 41). Other studies have shown that exposure to SO_2_ can result in an increase in neutrophils (33, 42, 45). Based on our data, neutrophils return to baseline levels by 1 month after chronic (x10) SO_2_ (125 ppm for 4 hours) exposure. Increased numbers of T lymphocytes are also present in Veterans with PDRS and have been observed in patients with idiopathic PAH (7, 46). Further studies will be required to determine whether adaptive immune activation contributes to the development of PH in this model.

Pulmonary vasculature abnormalities associated with SO_2_ inhalation have not been reported as an important component of lung pathology in previous studies (11, 47); however, high concentrations (200-850 ppm) of SO_2_ for 10 minutes has been shown to cause pulmonary vasoconstriction and an increase in pulmonary arterial pressure acutely in anesthetized dogs (48). Our study shows that pulmonary vascular pathology, including muscularization of the microvasculature, occurs as a long term consequence of repetitive SO_2_ exposure, at similar levels to those measured in other murine models of PH (49, 50). These data are consistent with human epidemiological studies demonstrating an association between SO_2_ exposure and increased morbidity and mortality from cardiovascular disease (51, 52). These findings are similar to those we have observed in lung biopsies from Veterans with PDRS (7). However, it remains unclear whether pathologic changes in the vasculature is associated with the development of PH in PDRS. Future clinical studies such as right heart catheterization will be required to definitively diagnosis PH in this population.

It is known that SO_2_ toxicity results from its ability to cause membrane damage, cell metabolism alterations, oxidative damage, DNA damage, alterations in signal transduction, and changes to ion channels (11, 53–56). The most extensively studied mechanism for SO_2_ toxicity is oxidative stress (9, 55, 56) and our data show excessive accumulation of superoxide after SO_2_ exposure in lung tissue, as well as pulmonary endothelial cells. Further, our studies show that ROS production after SO_2_ exposure is associated with increase in the lipid peroxidation, formation of isoLGs, Sirt3 inhibition, and reduction of SOD2 activity, as described in other pathological conditions, including hypertension (18, 57–59). In our studies, treatment with the isoLG scavenger 2-HOBA and Sirt3 overexpression rescued vascular remodeling after SO_2_ exposure. Notably, previous research has shown that increased ROS levels can cause mitochondrial oxidative stress, endothelial cell dysfunction, and vascular remodeling in PH (60). Furthermore, a recent clinical trial demonstrated that repeated oral administration of 2-HOBA is safe and well-tolerated, which indicates its potential for future clinical trials in cohort of Veterans with PDRS (61).

In summary, we developed a mouse model that reproduces many key features of human PDRS, including complex lung pathology and physiological abnormalities. We also identified PH as a long-term consequence of vascular remodeling in this model. We demonstrated involvement of the ROS-isoLGs-Sirt3-SOD2 pathway in the development of pulmonary vasculature pathology, thus shedding light on potential mechanisms relevant to human PDRS. Further studies in human PDRS should determine whether evidence of oxidative stress is present in the lungs of these patients and whether PH develops in these patients as a potential explanation for exertional dyspnea and exercise limitation.

## Acknowledgments

The authors thank David H. Wasserman and Louise Lantier in the Vanderbilt Mouse Metabolic Phenotyping Center for assistance with physical exercise testing.

## Author Contributions

SSG, RIS, DSN, and VVP conducted histopathological studies; BWR, SSG, and VVP developed model; JMRB, WH, and SSG performed flow cytometry; JJG and GUV contributed to collagen evaluation; DCN, JDW, and SSG conducted physiological studies; AED and GUV performed western blotting; SID, TSB, and VVP provided scientific and technical knowledge; SSG and TSB wrote the manuscript; VVP and TSB designed and co- supervised the project. The final version of the manuscript was approved by all authors.

